# Control strategies for the timing of intracellular events

**DOI:** 10.1101/714642

**Authors:** Mengfang Cao, Baohua Qiu, Jiajun Zhang, Tianshou Zhou

**Affiliations:** Key Laboratory of Computational Mathematics, Guangdong Province, P. R. China; School of Mathematics, Sun Yat-Sen University, Guangzhou, 510275, P. R. China

**Keywords:** first passage time, event timing, bursty gene expression, feedback regulation

## Abstract

While the timing of intracellular events is essential for many cellular processes, gene expression inside a cell can exhibit substantial cell-to-cell variability, raising the question of how cells ensure precision in the event timing despite such stochasticity. We address this question by analyzing a biologically reasonable model of gene expression in the context of first passage time (FPT), focusing on two experimentally measurable statistics: mean FPT (MFPT) and timing variability (TV). We show that: (1) transcriptional burst size (BS) and burst frequency (BF) can minimize the TV; (2) translational BS monotonically reduces the MFPT to a nonzero low bound and can minimize the TV; (3) the timescale of promoter kinetics can minimize both the MFPT and the TV, depending on the ratio of the off-switching rate over the on-switching rate; and (4) positive feedback regulation of any form can all minimize the TV, whereas negative feedback regulation of transcriptional BF or BS always enhances the TV. These control strategies can have broad implications for diverse cellular processes relying on precise temporal triggering of events.

## 1. Introduction

The timing of intracellular events is pivotal for many cellular processes, ranging from cellular responses to external stimuli to cell fate decision (1–8), cell differentiation (9–11), cell apoptosis (1, 12, 13), and cell cycle (14–18). For example, an activated gene may be required to reach in a precise time a threshold level of the regulatory protein expression that triggers a particular downstream signal pathway (16, 19, 20). Also for instance, fractional killing of a cell population (i.e., some cells in a clonal population die while others survive) depends on the time that time-keeping proteins reach threshold levels (called arrival time for brevity), e.g., cells must reach a threshold level of p53 to execute apoptosis and this threshold was experimentally shown to increase with time (21), and exposure of an isogenic bacterial population to a cidal antibiotic typically fails to eliminate a small fraction of refractory cells (22). Similarly, fractional killing of a cancer cell population by chemotherapy depends also on the arrival time (21, 23) the shorter the arrival time is, the more are the cells killed, indicating the therapeutic efficacy is better. In a word, timing events are ubiquitous. On the other hand, intracellular events take place often in a stochastic manner. This necessarily leads to variability in the event timing (which will be called timing variability in this paper). It is unclear how stochastic sources of timing events affect the timing of events. Characterization of control strategies for buffering timing variability is critically needed to understand reliable functioning of diverse intracellular pathways relying on precision in the timing as well as the efficacy of drugs depending on the time that drug-dependent regulatory proteins reach threshold levels.

Many of many cellular processes are based on gene expression, which is inherently noise due to low copy numbers. This stochasticity naturally gives rise to the cell-to-cell variability in the threshold-crossing time with potential consequences on biological functions and phenotypes and increasing experimental evidence has also shown timing variability and its consequences (20). However, stochastic sources of gene expression may be complex, since it would involve recruitment of transcription factors and polymerases (24–29), transitioning between active (on) and inactive (off) states of promoter (30,31), and chromatin remodeling (32–35). Each of these processes can all affect the crossing of threshold events. Given complexity of gene expression, two questions naturally arise: Are there optimal strategies to regulate the synthesis of a protein to ensure that an intracellular event will occur at a precise time while minimizing deviations or noise about the protein mean? How is the timing variability controlled in more realistic situations of gene expression?

Mathematically, threshold crossing can be formulated as a first passage time (FPT) problem (36). There have already been many works that used FPT frameworks to study the timing of events in biological and physical sciences (37–40) and the obtained results have provided insights into how model parameters shape statistical fluctuations in the event timing. Recently, Ghusinga, et al. (37) analyzed a simplified model of stochastic gene expression in the context of FPT. They claimed that for a stable long-lived protein, the optimal strategy is to express the protein at a constant rate without any feedback regulation, and any form of feedback (positive, negative, or any combination of them) will always amplify noise in event timing, whereas for an unstable protein, a positive feedback mechanism provides the highest precision in timing. Unfortunately, however, these qualitative results are not always correct but depend on the detail of feedback regulation (5, 8, 41–45). In other words, the strategies for control of the variability in the timing of intracellular events are not elucidated. This motivates the study of this paper.

In order to conclude biologically reasonable control strategies for timing variability, we use an experimentally validated and extensively used stochastic model of gene expression (46–49) to address the above questions. This model, an extended version of the common on-off model of gene expression (50–52), considers complexity of gene expression such as switching between promoter states, transcriptional and translational bursts as well as two kinds of feedback regulations with different forms (i.e., transcriptional regulations of burst size (BS) and burst frequency (BF) by positive or negative feedback). First, we formulate threshold crossing as a FPT problem where an event is triggered once a regulatory protein reaches a critical threshold, and establish a chemical master equation for this FPT problem. Second, we numerically solve this equation and analyze the FPT statistics. Third, we obtain qualitative results independent of the choice of model parameter values, e.g., both translational BS and positive feedback can minimize the TV; the timescale of promoter kinetics can minimize both the MFPT and the TV; and negative feedback always increases the TV. Our qualitative results actually give strategies for controlling timing variability, and can have broad implications for diverse cellular processes that rely on precise temporal triggering of events.

## 2. Stochastic model formulation and FPT calculation

### 2.1 Model description

Here, we simply describe an extended version of the common on-off model of stochastic gene expression, referring to Fig. 1(A). This model assumes that a gene promoter has one active (ON) and one inactive (OFF) states, between which there are transitions. Denote by *k*_on_ and *k*_off_ transition rates from OFF to ON and vice versa, respectively. Assume that the gene is initially at ON state at time *t* = 0 and begins to express a regulatory protein that a constant degradation rate denoted by δ. The intracellular event of interest is triggered once this protein reaches a threshold level in the cell. In addition, we assume translation in bursts and incorporates feedback regulation by considering ON- and/or OFF-switching rates as functions of the protein level, i.e., *k*_on_ (*x*) and *k*_off_ (*x*), where *x* represents the number of protein molecules, which is a function of time *t*, i.e., *x* = *x* (*t*). To explore the effects of feedback on the timing of events, we adapt Hill-type functions to describe regulation. Specifically, transcriptional burst frequency (BF) regulation and transcriptional burst size (BS) regulation are mathematically described as

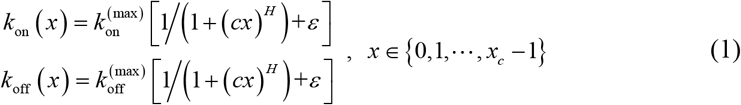

where 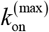 and 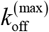 are the maximum transcription rates of *k*_on_and *k*_off_ respectively, *c* represents feedback strength, *ε* is a small positive constant describing the promoter leakiness, and *x*_*c*_ represents a fixed threshold. Note that Hill coefficient *H* = 0 represents no feedback, *H* > 0 and *H* < 0 represent negative and positive feedbacks respectively for transcriptional BS regulation whereas *H* > 0 and *H* < 0 represent psositive and negative feedbacks for transcriptional BF regulation.

**Figure1.**
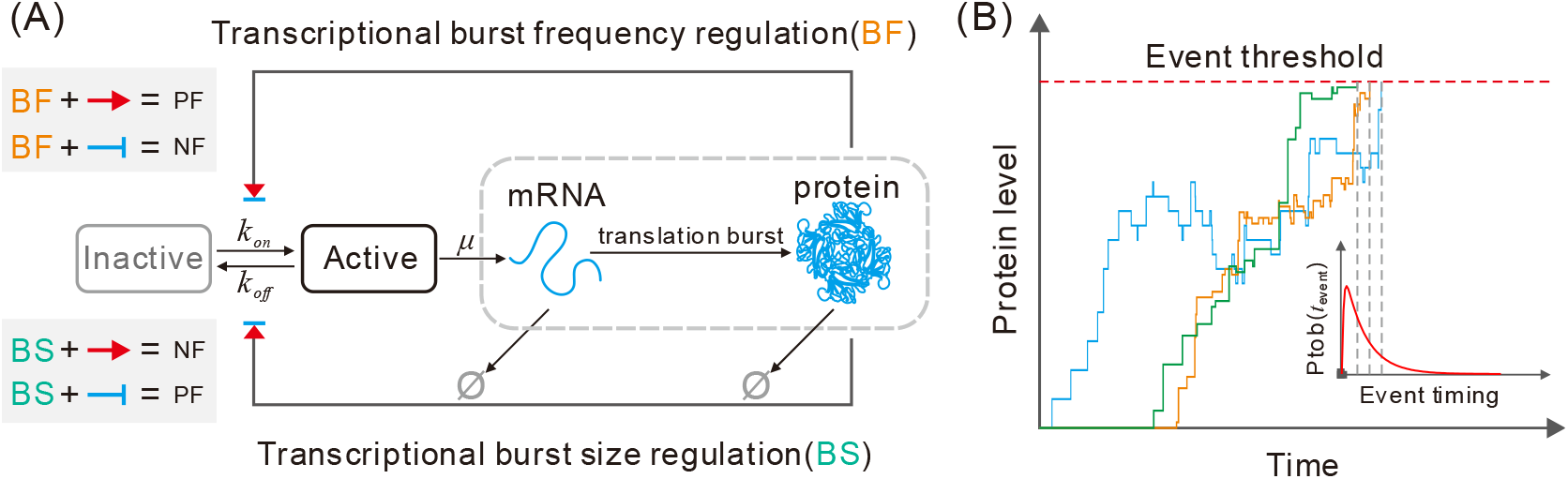
Modeling event timing as a first passage time (FPT) problem. (A) Schematic depiction of a gene model, where we assume that the gene switches between active and inactive states and when the gene is at active state, the DNA is transcribed into mRNAs with a constant rate, which are further translated into proteins. The resulting proteins regulate the switch rate from inactive to active states or vice versa or both, where BF and BS represent burst frequency and burst size, and PF and NF represent positive feedback and negative feedback. (B) Schematic description of the timing of an intracellular event that is formulated as the FPT for the protein level to reach a critical threshold, where the right-below inset shows a distribution of FPTs.

The translational burst approximation is based on assuming short-lived mRNAs, that is, each mRNA degrades instantaneously after producing a burst of *B* protein molecules. In agreement with experimental and theoretical studies (51, 52), *B* is assumed to follow a geometric distribution

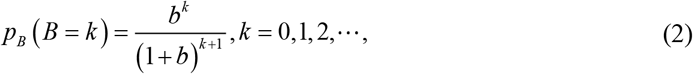

where *b* represents the mean protein burst size. Then, we can show *p*_*B*_ (*B* ≥ *k*) = *b*_*k*_/(1+*b*)^*k*^, *k* = 0,1,2, …. In what follows, we denote *p*_*B*≥*k*_ ≡ (*B* ≥ *k*) for simplicity.

For convenience, let *s* = 0 and *s* = 1 represent inactive and active states, respectively. Denote by *p*_*s*_ (*t*; *x*) the probability that the protein has *x* molecules at state *s* at time *t* and stipulate *p*_*s*_ ≡ 0 if *x* < 0, where *s* = 0 or 1. Under the above settings or assumptions, the chemical master equation for the time evolution for this probability can be then described as

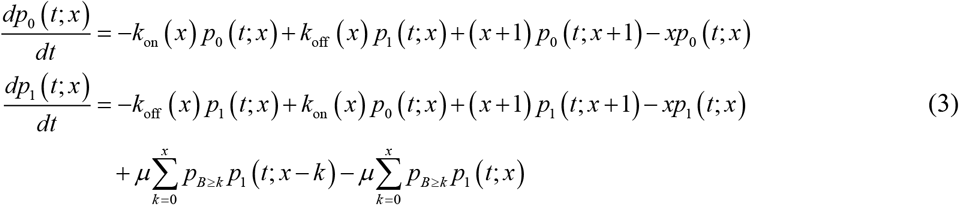

where *x* = 0,1, …, *x*_*c*_ −1. Denote by *p* (*t*; *x*) the total probability that the protein has *x* molecules at time *t*. Then, *p* (*t*; *x*) = *p*_0_ (*t*; *x*) + *p*_1_(*t*; *x*).

On the other hand, the time to an event is the FPT for *x*(*t*) to reach a threshold *x*_*c*_ starting from a zero initial condition *x* (0) = 0 for the first time, referring to Fig. 1(B). It is mathematically described by the following random variable

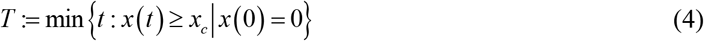

and can be interpreted as the time taken by a random walker to reach a defined point for the first time. Denote by *f*_*T*_(*t*) the probability density function (PDF) of the FPT. The left problem is how we give *f*_*T*_(*t*) based on Eq. (4).

### 2.2 Distribution of the FPT

Let ***p***(*t*; *x*) = [*x*_0_(*t*; *x*), *p*_1_(*t*; *x*)]^T^ be a two-dimensional vector, where T represents transpose. Since *x* takes finite values in Eq. (3), we enumerate all possible cases into a vector of the form:

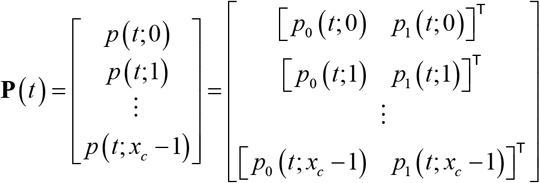

Assume that the initial vector of probabilities is **P**(**0**) = [[1,0],[0,0],…,[0,0]]^T^. Note that Eq. (4) can be rewritten as the following vector

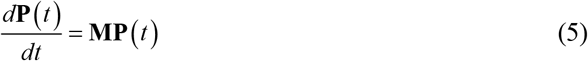

where the matrix **M** takes the form

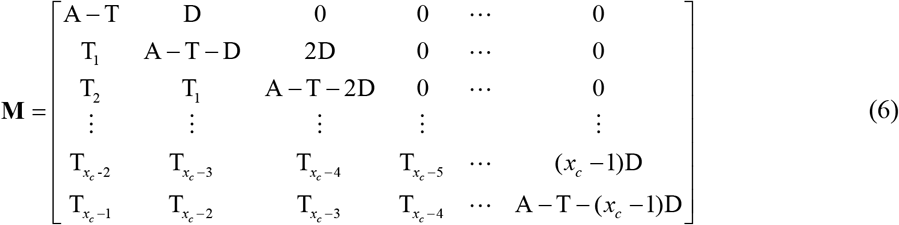

Here, matrix A describes state transitions, matrices T and T_*j*_ with 1 ≤ *j* ≤ *x*_*c*_ −1 are associated with burst transcription, and matrix D describes degradation. These matrices take the following forms

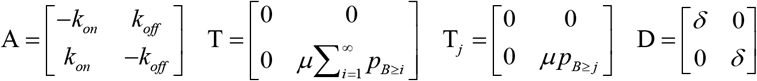

Note that Eq. (4) allows an analytical solution of the form

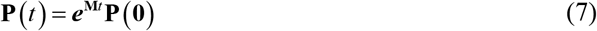

Next, we give the formal expression of the PDF of FPT, *f*_*T*_(*t*). For this, we introduce vector 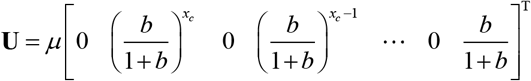. According to the definition of FPT, we then know that the PDF of FPT is given by:

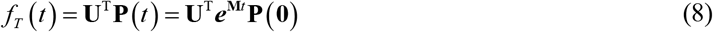

### 2.3 Moments of the FPT

Although the PDF of FPT given by Eq. (8) provides complete characterization of the event timing, we are particularly interested in the lower-order statistical moments of FPT. Here, we exploit the structure of matrix **A** to obtain analytical formulas for the first and second-order moments of FPT.

First, the *n* -order raw moment of FPT is calculated according to

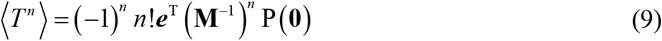

where ***e*** = [1,…,1]^T^ is a *n* -dimensional column vector.

Second, the mean FPT (MFPT), i.e., the first order moment of FPT is calculated according to

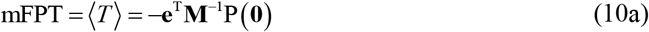

The TV, which also represents the noise intensity of FPT, is measured by the square of the coefficient of variation according to

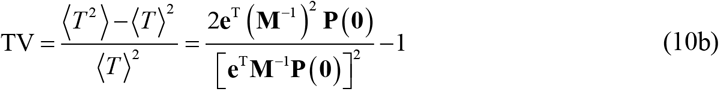

The above formulae give only the formal expressions of statistical quantities for FPT, and their practical calculations (in particular, the one of inverse matrix **M**^−1^) need to seek numerical methods. In order to obtain analytical results of MFPT and TV, we consider the limit of large mean burst size (*b*). In this case, we can show (seeing Appendix for derivation)

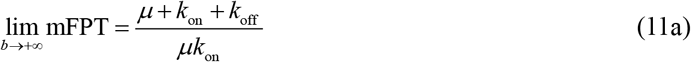

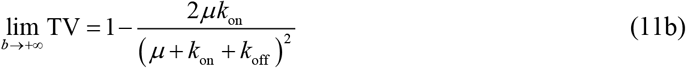

These analytical results have been numerically verified (see the section of main results).

Finally in this subsection, we point out that to model the complexity of gene expression mentioned above, we let model parameters change in broad yet biologically reasonable ranges.

## 3. Main results

### 3.1 Optimal burst control strategy

Bursty expression is a common way of gene expression. Our model involves two kinds of bursts: transcriptional burst and translational burst. For the former, BS is *μ*/*k*_off_ and BF is *k*_on_ (if we set δ= 1) in the case of no feedback (corresponding to *H* = 0). For the latter, the mean BS, *b* has been assumed to follow a geometric distribution given by Eq. (2). Here, we examine how these bursts influence the event timing (focusing on MFPT and TV) in the case of no feedback. Numerical results are shown in Fig. 2.

**Figure 2.**
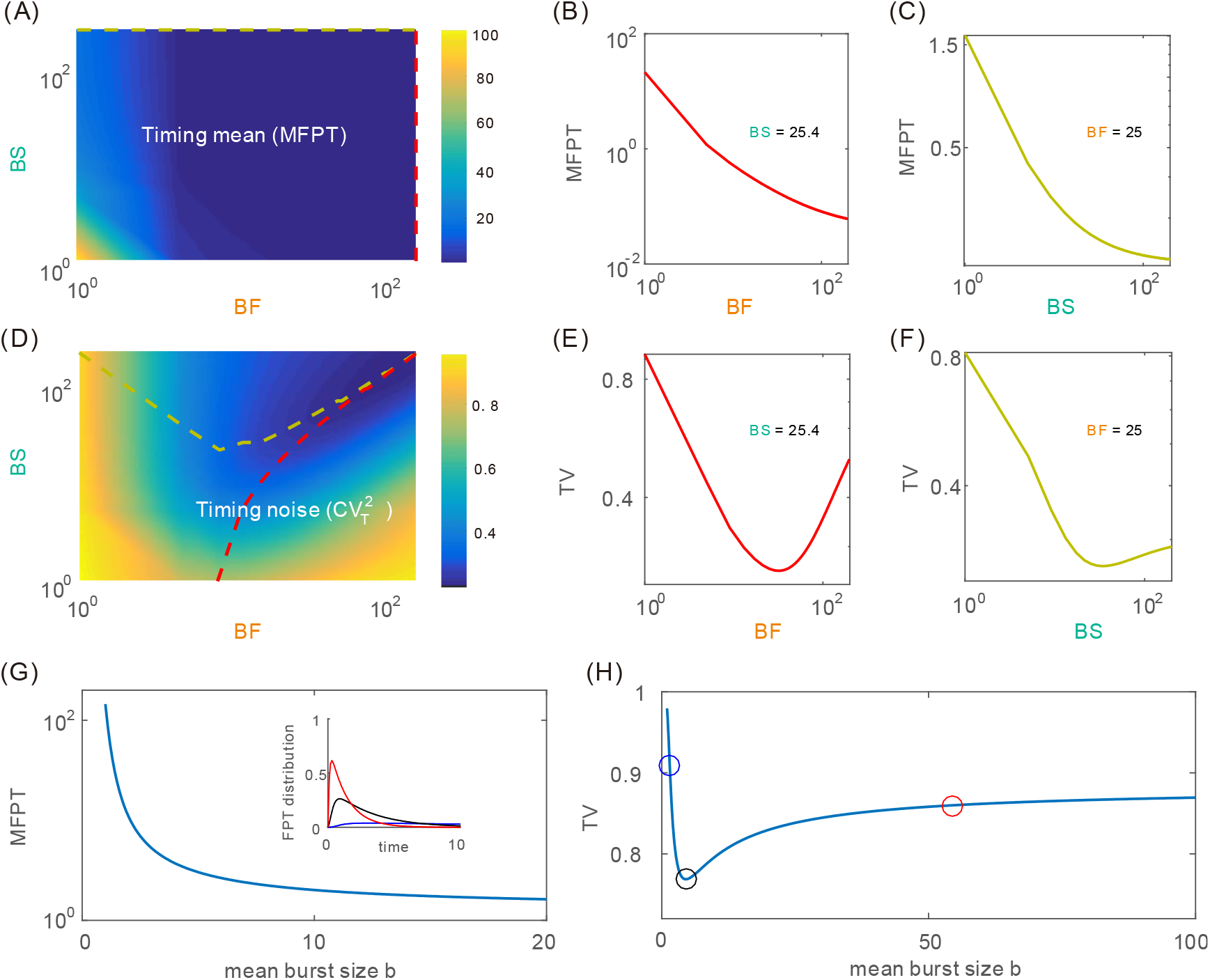
Influences of bursts on the event timing in the case of no feedback. (A) Mean first passage (MFPT) as a function of transcriptional burst size (BS) and transcriptional burst frequency (BF), where canary dashed and red dashed lines represent the paths of the minimum values of MFPT under fixing BF and BS, respectively, and parameter values are set as *x*_*c*_ = 20, *b* = 3, *k*_off_ = 2, *μ* : 2 ~ 400 and *k*_on_:1 ~ 200. (B,C) Special cases of (A), where (B) shows dependence of MFPT on BF for a fixed BS and (C) shows dependence of MFPT on BS for a fixed BF. (D) Timing variability (TV) as a function of BS and BF, where canary dashed and red dashed lines represent the paths of the minimum values of TV under fixing BF and BS, respectively, and parameters values are set as in (A). (E,F) Special cases of (D), where (E) shows dependence of TV on BF for a fixed BS and (F) shows dependence of MFPT on BS for a fixed BF. (G) MFPT as a function of mean translational BS, where the inset shows three distributions of FPT. (H) TV as a function of mean translational BS. In (G) and (H), parameter values are set as *x*_*c*_ = 20, *k*_on_ = 1, *k*_off_ = 2, and *μ* = 10.

Figure 2(A) shows a global scenario for the dependence of MFPT on both transcriptional BS and transcriptional BF. From this panel, we observe that the MFPT is a monotonically decreasing function of BF for a fixed BS (referring to Fig. 2(B)) and of BS for a fixed BF (referring to Fig. 2(C)). This qualitative result is in accordance with our intuition and is not strange. Figure 2(D) also shows a global scenario for the dependence of TV on both BS and BF. From this panel, we observe that the curve of TV vs transcriptional BF for a fixed transcriptional BS is convex (referring to Fig. 2(E)), implying that there is an optimal transcriptional BF such that the TV is lowest or that transcriptional BF can minimize the variability in the event timing. For a fixed transcriptional BF, there is a critical value of transcriptional BS such that TV reaches an optimal value, referring to Fig. 2(F).

Figures 2(G) and 2(H) show the dependences of MFPT and TV on mean translational burst size (*b*), respectively. We observe from Fig. 2 (G) that the MFPT is a monotonically decreasing function of *b*, implying that translational burst reduces the mean time that the protein reaches a given threshold. In addition, as *b* tends to infinity, the MFPT has a finite, positive limit (i.e., the MFPT eventually tends to a stable value) given analytically by Eq. (11a). The inset in Fig. 2(G) demonstrates three different distributions of FPT. An interesting phenomenon is shown in Fig. 2(H), from which we observe that there is an optimal mean translational burst size such that the variability in the event timing is lowest. Moreover, there is a finite, positive limit as *b* tends to infinity, seeing the analytical expression given by Eq. (11b).

The results shown in Fig. 2 indicate that transcriptional and translational bursts both are an effective mechanism of controlling the timing of intracellular events.

### 3.2 Optimal promoter-switching control strategy

In general, the switching rates between promoter states are not fixed constants but are regulated often by, e.g., external signals. Experimental evidence has pointed to the fact that promoter fluctuations generated due to stochastic switching between promoter states are a major source of cell-to-cell variability (i.e., gene expression noise). Note that in our model, two switching parameters, *k*_on_ and *k*_off_, can characterize promoter kinetics. In this subsection, we allow these two parameters to change in biologically reasonable ranges and consider only the case of *H* = 0 (i.e., without self-feedback). In order to characterize the size of promoter fluctuations, we introduce a common timescale factor (denoted by *α*) for *k*_on_ and *k*_off_ while keeping the ratio of *k*_on_ over *k*_off_ fixed. Apparently, this factor has the following property: the smaller the ratio is, the more slowly does the promoter switch between two states, implying that promoter fluctuations are larger, and the larger the ratio is, the more quickly does the promoter switch between two states, implying that promoter fluctuations are smaller. Here, we are interested in how the timescale factor (*α*) of promoter kinetics impacts the mean of FPT (i.e., MFPT) and the variability in the event timing (i.e., TV). For clarity, we consider two cases: (1) *k*_on_/*k*_off_ is fixed but *α* changes; (2) *α* is fixed but *k*_on_/*k*_off_ changes. Numerical results are demonstrated in Fig. 3.

**Figure 3.**
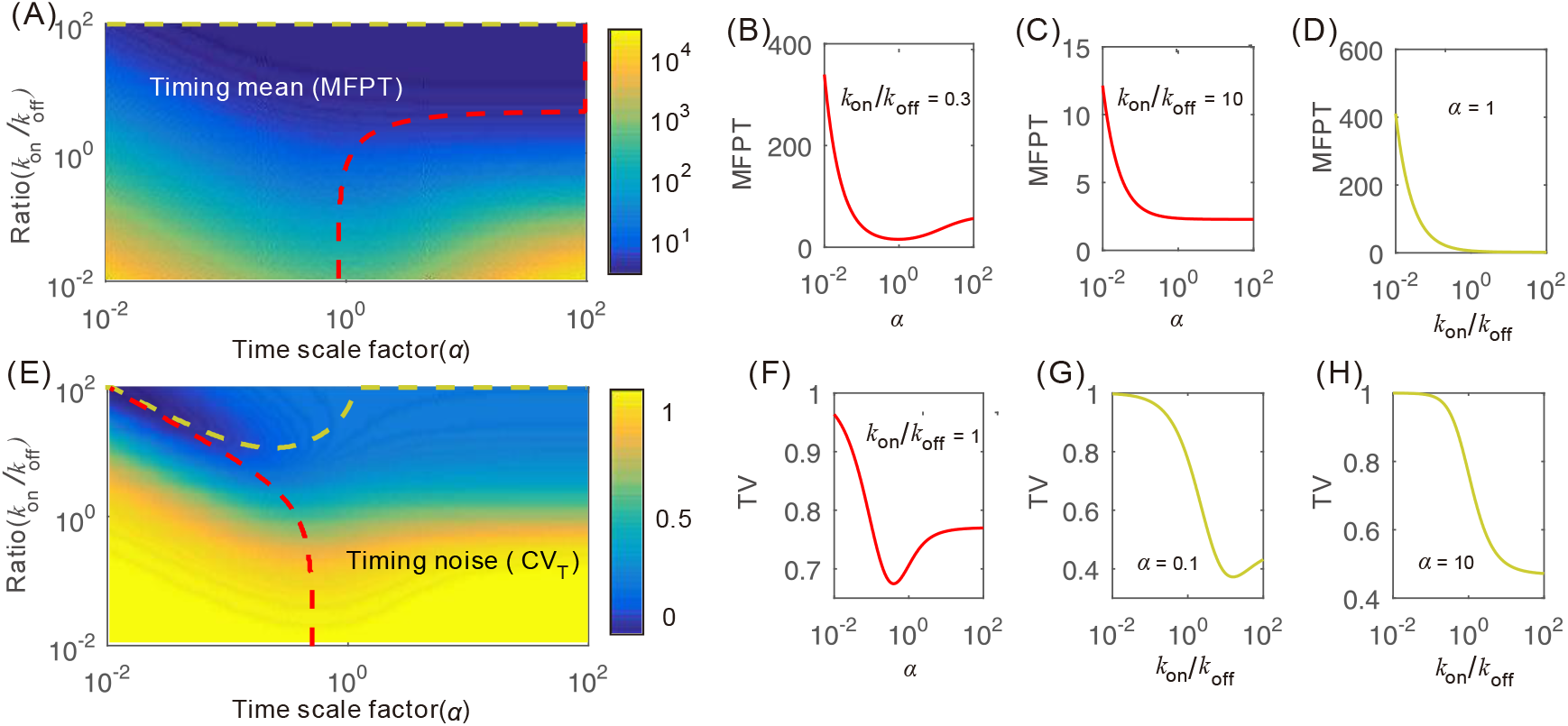
Influence of promoter kinetics on the event timing in the case of no feedback. (A) The global scenario for dependence of MFPT on both timescale factor *α* and ratio *k*_on_/*k*_off_, where canary dashed and red dashed lines represent the paths of the minimum value of MFPT under fixing *α* and *k*_on_/*k*_off_, respectively. (B-D) Special cases of (A), where parameter values are set as *x*_*c*_ = 20, *b* = 2, *k*_off_ = 1, *δ* = 1 and *μ* = 10. (E) The global scenario for dependence of the TV on both time scale factor *α* and ratio *k*_on_/*k*_off_, where parameter values are set as in (A).

In order to show the global scenario of how timescale factor, *α*, and the ratio of off-switching rate over on-switching rate, *k*_on_/*k*_off_, altogether affect the time that the protein of intracellular events reaches a threshold for the first time, we plot Fig. 3(A), a three-dimensional pseudo-diagram, where a canary dashed line and a red dashed line represent the paths of the minimum value of MFPT under fixing *α* and *k*_on_/*k*_off_, respectively. We observe that if ratio *k*_on_/*k*_off_ is fixed and small than 1, then timescale factor *α* can minimize the mean first passage time or there is an optimal timescale factor such that the MFPT is minimal, referring to Fig. 3(B). If ratio *k*_on_/*k*_off_ is fixed and larger than 1, then the MFPT is a monotonically decreasing function of timescale factor *α* (referring to Fig. 3(C)), implying that the timescale of promoter kinetics can enhance response. Figure 3(D) shows that the MFPT is also a monotonically decreasing function of ratio *k*_on_/*k*_off_ in the case of *α* = 1, implying that the ratio of the off-switching rate over the on-switching rate can also enhance response.

Figure 3(E) shows the global scenario of how timescale factor, *α*, and the ratio of on-switching rate over off-switching rate, *k*_on_/*k*_off_, altogether affect the variability in the event timing. Two dashed curved indicated in this figure represent two special paths of the minimum value of MFPT under the condition that *α* and *k*_on_/*k*_off_ are fixed, respectively. Figure 3(F-H) shows results in special cases of *α* and *k*_on_/*k*_off_. Specifically, if ratio *k*_on_/*k*_off_ is fixed and small than or equal to 1, then timescale factor *α* can minimize the variability in the event timing or there is an optimal timescale factor such that the TV is minimal, referring to Fig. 3(F). If ratio *α* is fixed and smaller than 1, then the ratio of the off-switching rate over the on-switching rate can also minimize the TV, referring to Fig. 3(G). If ratio *α* is fixed and larger than 1, then the TV is a monotonically decreasing function of ratio *k*_on_/*k*_off_, referring to Fig. 3(H).

### 3.3 Optimal feedback control strategy

Our model introduces two kinds of feedbacks: the one is to regulate the transition rate from off to on states and the other to regulate the transition rate from on to off states, referring to Fig. 1(A). These feedbacks may be positive or negative. Here, the question we are interested in is how feedback strength represented by *c* affects the timing of intracellular events (in fact, the expression level of the protein). We may assume that *c* changes in the interval of 10^−4^ ~ 1 due to our setting. In order to find the optimal feedback mechanism, our strategy is to change 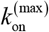 or 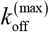 in Eq. (1) while other parameters are kept fixed. Numerical results are demonstrated in Fig. 4, which shows all possible modes of the curve for the dependence of timing variability on feedback strength.

**Figure 4.**
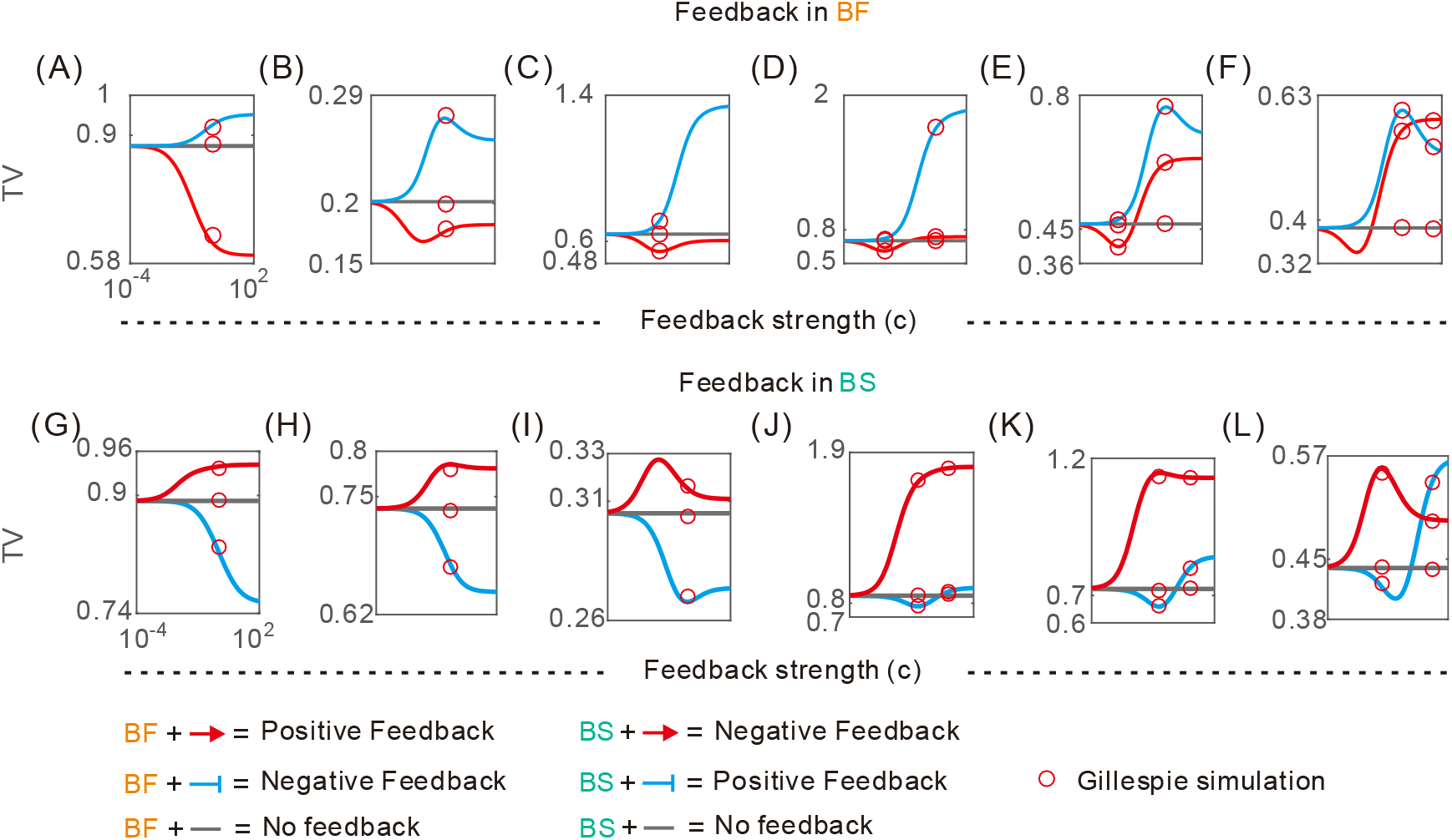
Influence of feedback on the event timing. , where the MFPT is fixed at 10. (A-F) TV is plotted as a function of feedback strength *c* for different regulation strategies of transcriptional BF, where parameters values are set in the table1. (G-L) TV is plotted as a function of *c* for different regulation strategies of transcriptional BS, where parameters values are set in the table 2. In all subfigures, red and blue curves correspond respectively to positive and negative feedbacks, the grey lines correspond to no feedback, and red circles correspond to the results obtained by the Gillespie algorithm. positive feedback. *H* = 1 for negative feedback, *H* = 0 without feedback, and *H* = −1 for positive feedback.

First, we examine the case of transcriptional BF regulation. Numerical results are demonstrated in Fig. 4(A-F). These panels show six modes for the dependence curve of timing variability (TV) vs feedback strength (*c*). In all the modes, the curve for TV vs *c* in the case of negative feedback is beyond that in the case of no feedback but may be below (referring to Fig. 4(A-C)) or cross (referring to Fig. 4(D-F)) that in the case of no feedback. Specifically, Fig. 4(A) shows that the curve for TV vs *c* is monotonically increasing *c* in the case of negative feedback but monotonically increasing function of *c* in the case of positive feedback and below that in the case of no feedback. Fig. 4(B) shows that in each of feedback regulation cases, the curve for TV vs *c* is not monotonically increasing *c* but there is a critical feedback strength such that TV reaches an optimal value (the maximum for negative feedback but the minimum for positive feedback). Different from Fig. 4(B), Fig. 4(C) shows that TV does not has an optimal value in the case of negative feedback but has an optimal value in the case of positive feedback. The only difference between Fig. 4(C) and Fig. 4(D) is that the curve for TV vs *c* in the case of positive feedback crosses that in the case of no feedback. Fig. 4(E) is similar to Fig. 4(D) but TV has an optimal value in both regulation cases. The only difference between Fig. 4(F) and Fig. 4(E) is that the curve for TV vs *c* in the case of positive feedback intersects with that in the case of no feedback.

Then, let us examine the case of transcriptional BS. Numerical results are demonstrated in Fig. 4(G-L). From theses panels, we observe that there are also six modes for the dependence curve of TV vs *c*. Overall, these modes are similar to those in Fig. 4(A-F). The detailed description is omitted.

We point out that all results shown in Fig. 4 are obtained in the case of Hill coefficient *H* = 1. However, *H* may take other values. Fig. S1 in Appendix demonstrates numerical results obtained in the case of *H* ≠ 1, showing that Hill coefficient can also influence the mode for the dependence curve of TV as *c*. In addition, we point out that the qualitative results claimed in ref. (37) are not always correct but depend on details of feedback regulation.

## 4. Discussions

Previous studies focused on cell-to-cell variability or fluctuations in the molecule numbers of gene products (53, 54), and less considered and event ignored the cell-to-cell variability in the timing of intracellular events. In this paper, using an experimentally validated and commonly used stochastic model of gene expression, we have systematically investigated how intracellular events cross critical thresholds, focusing on control strategies of MFPT and TV. We have demonstrated that the timescale of promoter kinetics can minimize both the MFPT and the TV, depending on the ratio of the off-switching rate over the on-switching rate; in contrast to negative feedback that always increases the TV, positive feedback can minimize the TV, independent of regulation forms; and translational BS monotonically reduces the MFPT to a nonzero low bound, whereas translational BF can minimize the TV.

Our model considered only the case of fixed thresholds. However, dynamically fluctuating thresholds are ubiquitous in biological regulatory systems. Such a kind of threshold has a strong biological background and is ubiquitous in biological regulatory systems. For example, consider a representative activity function of Hill type

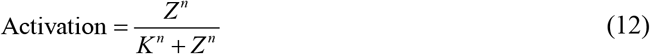

where *n* is a Hill coefficient, and *K* = *k*_−_/*k*_+_ represents a threshold (in fact a dissociation constant) of variable *Z*. In general, reaction rate *k*_+_ or *k*_−_ is regulated by external signals that are stochastically generated due to biochemical reactions, e.g., 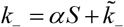, where *S*, an external signal, is stochastically generated, *α* and 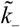 are positive constants. In this case, *K* is a dynamically fluctuating threshold of *Z*. This naturally raises questions: How does a dynamically fluctuating threshold impact event timing including first passage time and timing variability; What advantages does this kind of threshold in contrast to a fixed threshold? These questions are worth investigation.

We have seen that the noise in gene expression levels can affect both first passage time and timing variability and there are control strategies for buffering the noise in event timing. However, sources of gene expression noise may be complex, e.g., promoter noise generated by switching between multi-states of the promoter (55), and the noise resulting from chromatin remodeling, DNA methylation, and nucleosome positioning (56). These stochastic sources can influence not only gene expression levels but also threshold crossing. How they affect first passage time and timing variability remains unexplored.

Finally, the qualitative results obtained here can have broad implications for diverse cellular processes (as mentioned in the introduction) that rely on precise temporal triggering of events. In addition, fractional killing, a phenomenon occurring in, e.g., bacteria and drug therapy, relies on the time that relevant proteins reach threshold levels (called arrival time for brevity), e.g., exposure of an isogenic bacterial population to a cidal antibiotic typically fails to eliminate a small fraction of refractory cells (57), and cells must reach a threshold level of p53 to execute apoptosis and this threshold increases with time (58). Similarly, fractional killing of a cancer cell population by chemotherapy depends on the arrival time: the shorter the arrival time is, the more are the cells killed, indicating the therapeutic efficacy is better. Revealing the mechanisms behind these phenomena is significant and has potential application perspectives.

## Acknowledgments

This work was supported by grants 91530320, 11775314, 11475273, and 11631005 from the National Natural Science Foundation of China; 2014CB964703 from Science and Technology Department of China; 201707010117 from the Science and Technology Program of Guangzhou.

## APPENDIX

## A. Derivation of Eq. (11)

If burst size is large enough, then we can view that the regulatory protein generated by one-step transcription can cross a given threshold. In this case, we have the following model of gene expression

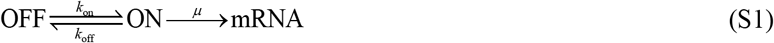

The corresponding chemical master equation takes the form

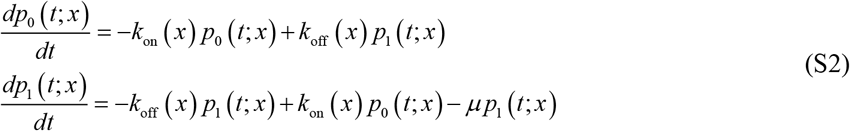

Using the denotations in the main text, we have

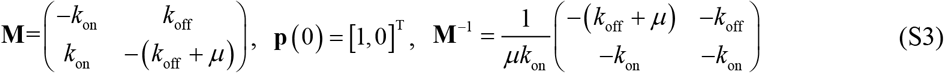

Thus, we can obtain the bounds of mean FPT, second-order moment of FPT, and the intensity of noise in event timing in the limit of large mean translational burst size as follows

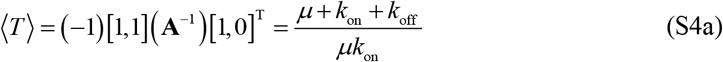

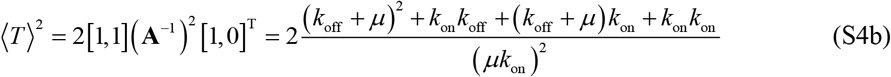

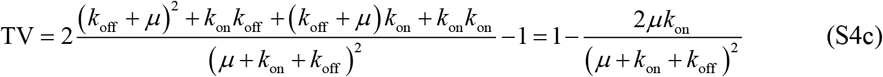

## B. Setting of parameter values

**Table 1.**
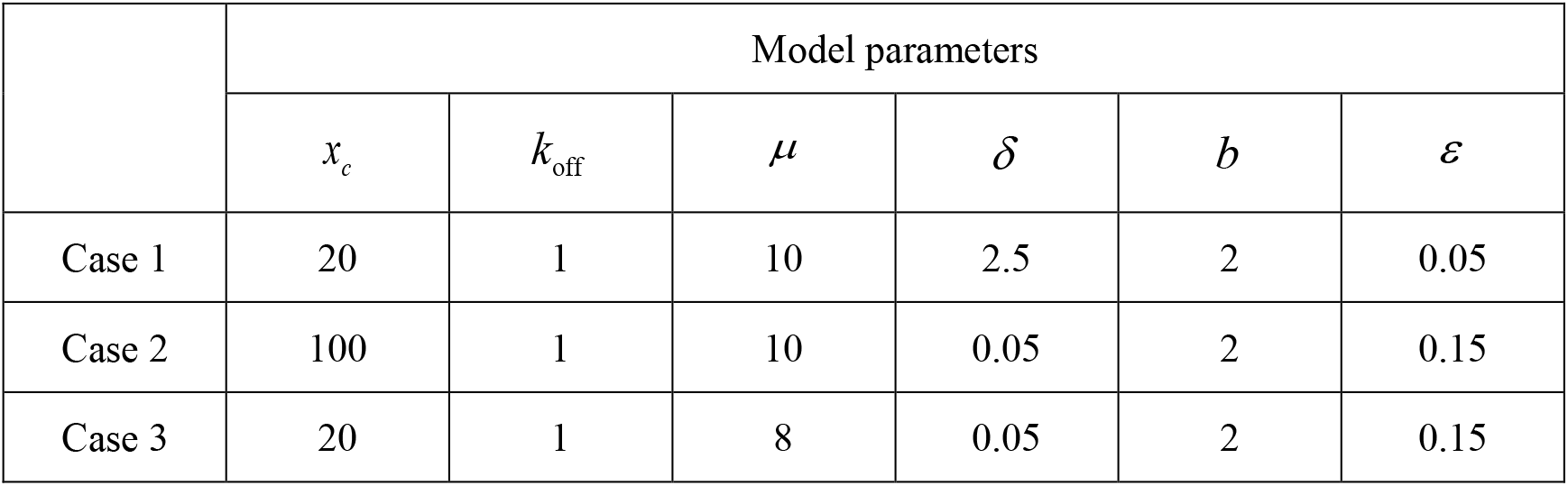
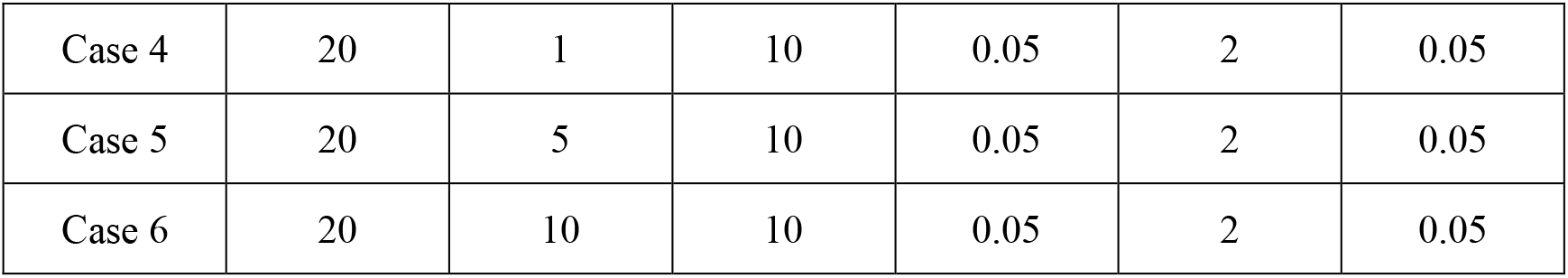
Setting of parameter values in the case of BS regulation

|  | Model parameters |  |  |  |  |  |
| --- | --- | --- | --- | --- | --- | --- |
| | $x_c$ | $k_{\text{off}}$ | $\mu$ | $\delta$ | $b$ | $\varepsilon$ |
| Case 1 | 20 | 1 | 10 | 2.5 | 2 | 0.05 |
| Case 2 | 100 | 1 | 10 | 0.05 | 2 | 0.15 |
| Case 3 | 20 | 1 | 8 | 0.05 | 2 | 0.15 |
| Case 4 | 20 | 1 | 10 | 0.05 | 2 | 0.05 |
| Case 5 | 20 | 5 | 10 | 0.05 | 2 | 0.05 |
| Case 6 | 20 | 10 | 10 | 0.05 | 2 | 0.05 |

**Table 2.** Setting of parameter values in the case of BS regulation

|  | Model parameters |  |  |  |  |  |
| --- | --- | --- | --- | --- | --- | --- |
| | $x_c$ | $k_{on}$ | $\mu$ | $\delta$ | $b$ | $\varepsilon$ |
| Case 1 | 20 | 1 | 10 | 1 | 3 | 0.05 |
| Case 2 | 20 | 1 | 10 | 0.35 | 2 | 0.5 |
| Case 3 | 80 | 1 | 10 | 0.05 | 2 | 0.05 |
| Case 4 | 20 | 1 | 10 | 0.05 | 18 | 0.05 |
| Case 5 | 20 | 1 | 10 | 0.05 | 9 | 0.05 |
| Case 6 | 20 | 1 | 10 | 0.05 | 2 | 0.05 |

**Table 3.** Parameter values for the different feedback results

|  | Feedback in BF Parameter Values |  |  |  |  |  |
| --- | --- | --- | --- | --- | --- | --- |
| | $x_c$ | $k_{off}$ | $\mu$ | $\delta$ | $b$ | $\varepsilon$ |
| Case1 | 50 | 1 | 6 | 0 | 2 | 0.15 |
| Case2 | 20 | 8 | 10 | 0 | 4 | 0.35 |
| Case3 | 20 | 10 | 10 | 0 | 2 | 0.001 |

**Table 4.** Parameter values for the different feedback results

|  | Feedback in BS Parameter Values |  |  |  |  |  |
| --- | --- | --- | --- | --- | --- | --- |
| | $x_c$ | $k_{on}$ | $\mu$ | $\delta$ | $b$ | $\varepsilon$ |
| Case1 | 20 | 5 | 10 | 0 | 8 | 0.5 |
| Case2 | 40 | 3 | 2 | 0 | 25 | 0.01 |
| Case3 | 20 | 6 | 15 | 0 | 2 | 0.01 |

## C. Supplementary figure: Effect of Hill coefficient

In the main text, we fix Hill coefficient at *H* = 1. However, *H* may take other values in some realistic cases. Here we address the question of how the curve for the dependence of timing variability on feedback strength changes for different values of *H* if other parameter values are kept fixed. Numerical results are demonstrated in Fig. S1, which shows several different modes. This figure indicates that the qualitative results in ref. (37) are not always correct.

**Figure S1.**
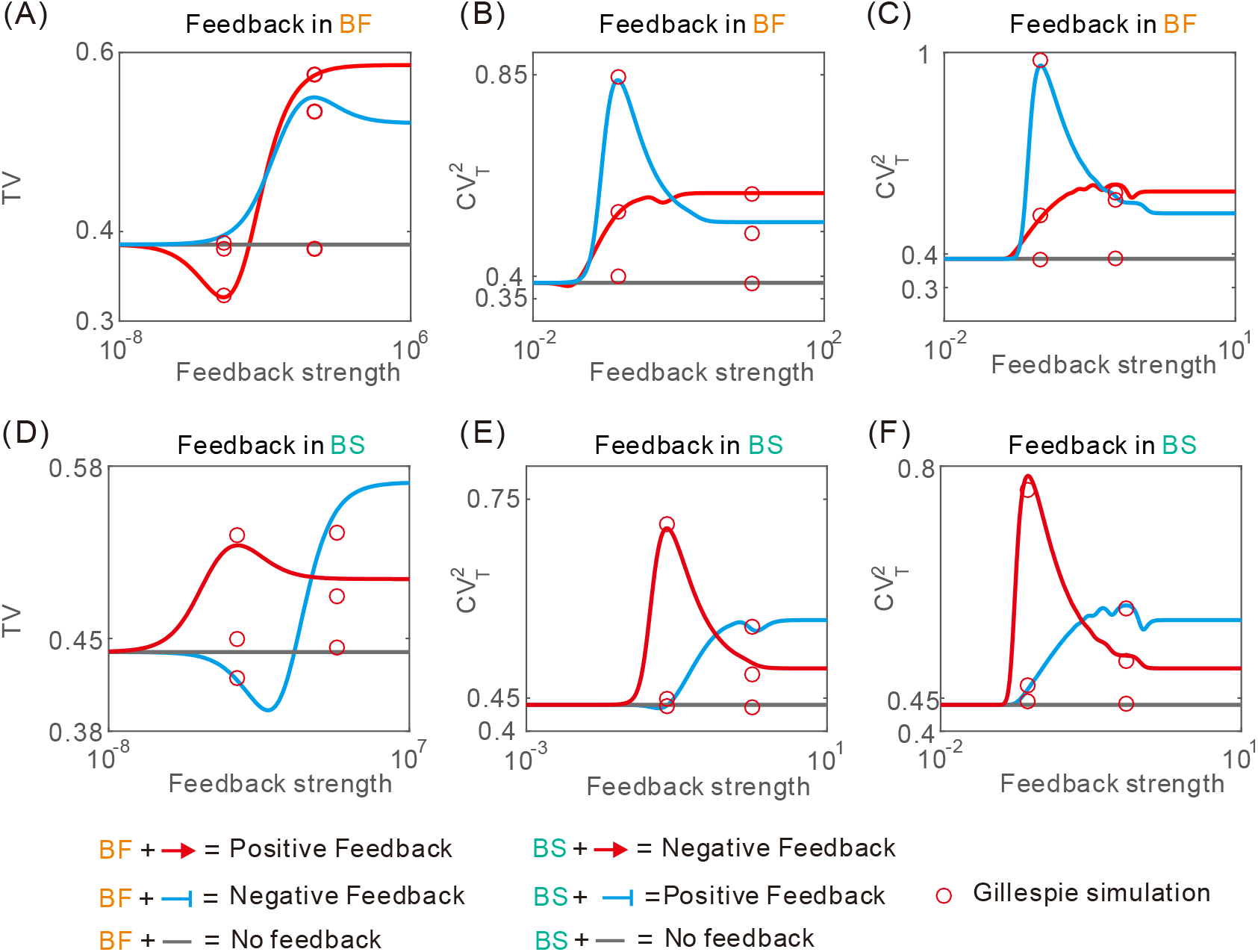
Influence of Hill coefficient on timing varibability. , where the MFPT is fixed at 10, and some parameter values are set as *x*_*c*_ = 20, *μ* = 10, *δ* 0.05, *k*_off_ = 10, *b* = 2, and *r* = 0.05. Red and blue curves correspond respectively to positive and negative feedbacks, grey lines correspond to no feedback and red circles correspond to the Gillespie algorithm. (A) TV is plotted as a function of feedback strength *c* for different regulation strategies of transcriptional BF, where negative feedback: *H* = 0.5, without feedback: *H* = 0, and positive feedback: *H* = −0.5. (B) Description is the same as (A), but negative feedback: *H* = 5, and positive feedback: *H* = −5. (C) Description is the same as (A), but negative feedback: *H* = 15, and positive feedback: *H* = −15. (D) TV as a function of *c* for different regulation strategies of transcriptional BS, where negative feedback: *H* = −0.5, without feedback: *H* = 0, and positive feedback: *H* = 0.5. (E) Description is the same as (D), but negative feedback: *H* = −5, without feedback: *H* = 0, and positive feedback: *H* = 5. (F) Description is the same as (D), but negative feedback: *H* = −15, without feedback: *H* = 0, and positive feedback: *H* = 15 .

We have explored optimal feedback for an unstable protein. Here we address the question of how the curve for the dependence of timing variability on feedback strength changes for a stable protein. Numerical results are demonstrated in Fig. S2.

**Figure S2.**
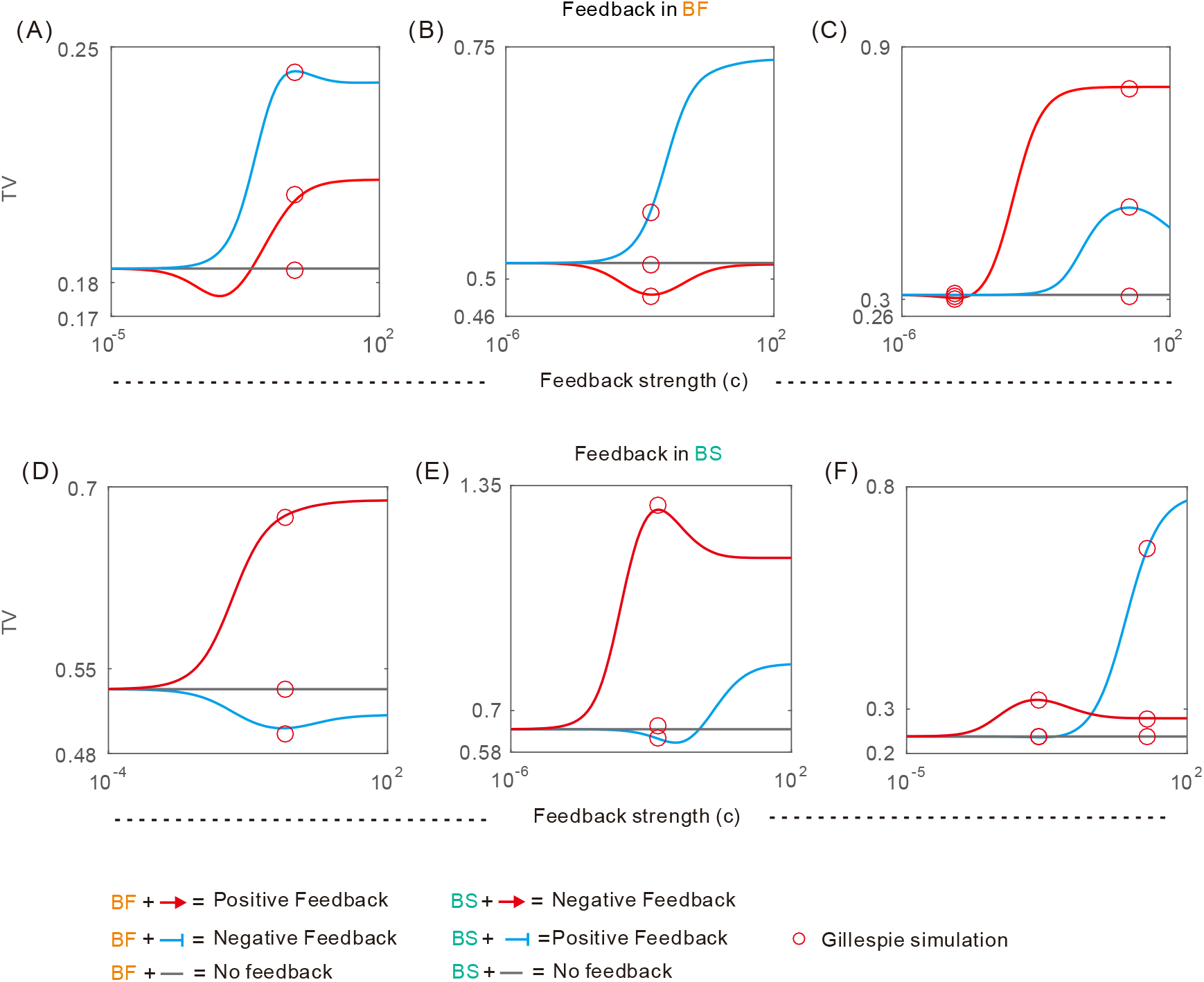
Influence of feedback for a Stable Protein. , where the MFPT is fixed at 10. (A-C) TV is plotted as a function of feedback strength *c* for different regulation strategies of transcriptional BF, where parameters values are set in the table 3. Red and blue curves correspond respectively to positive and negative feedbacks. *H* = 1 for positive feedback, *H* = 0 without feedback, and *H* = −1 for negative feedback. (D-F) TV is plotted as a function of *c* for different regulation strategies of transcriptional BS, where parameters values are set in the table 4. Blue and red curves correspond respectively to positive and negative feedbacks. In all subfigures, the grey lines correspond to no feedback, and red circles correspond to the results obtained by the Gillespie algorithm. feedback. *H* = 1 for negative feedback, *H* = 0 without feedback, and *H* = −1 for positive feedback.

From Fig. S2, we find that in the case of stable protein or unstable protein, the influence of feedback is similar.

In Fig. 3, we are interested in how the timescale factor (*α*) of promoter kinetics impacts the mean of FPT (i.e., MFPT) and the variability in the event timing (i.e., TV), here, we further consider the situation of stable proteins, how they affect MFPT and TV.

**Figure S3.**
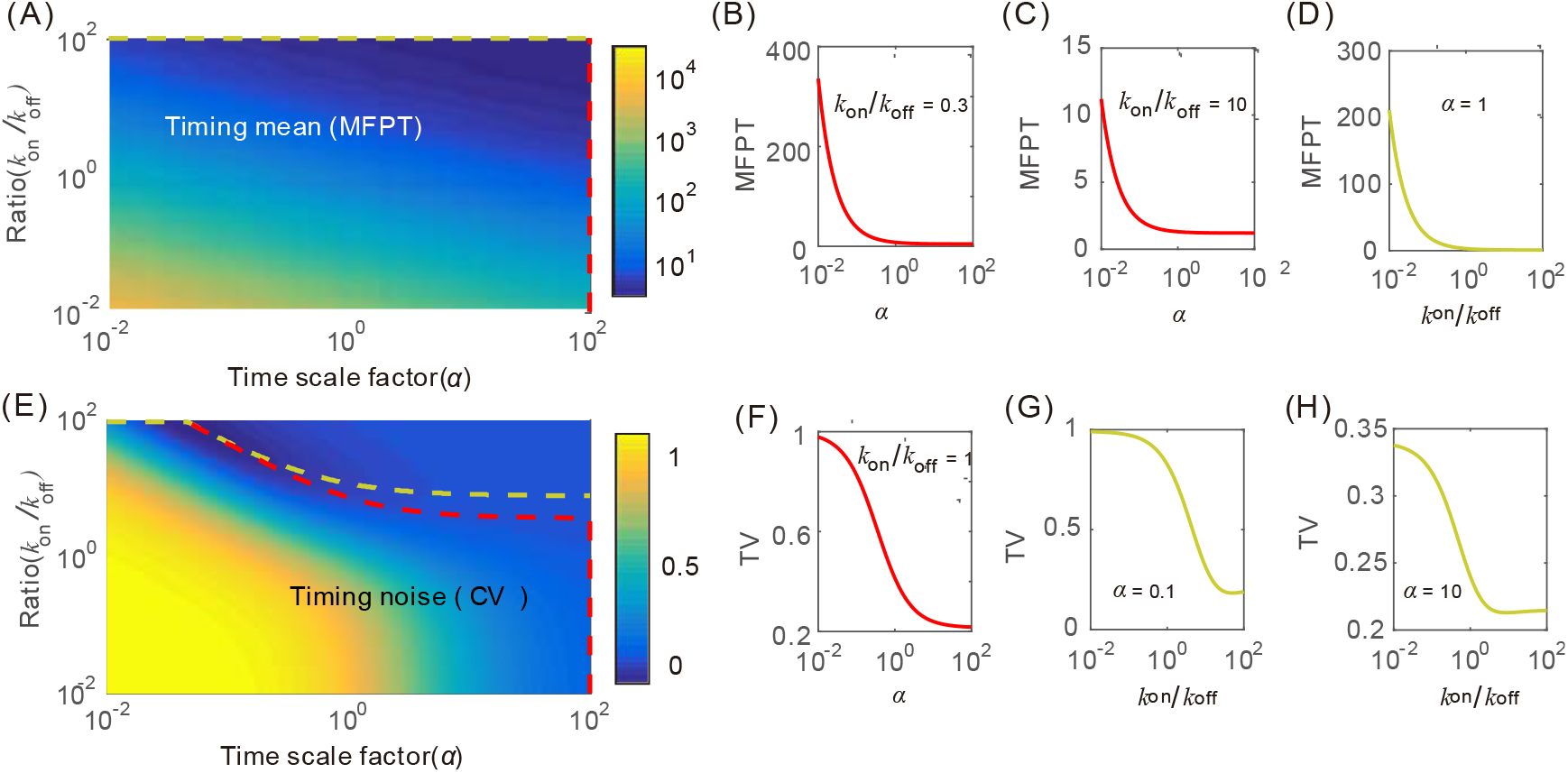
Influence of promoter kinetics on the event timing in the case of no feedback and stable protein. (A) The global scenario for dependence of MFPT on both timescale factor *α* and ratio *k*_on_/*k*_off_, where canary dashed and red dashed lines represent the paths of the minimum value of MFPT under fixing *α* and *k*_on_/*k*_off_, respectively. (B-D) Special cases of (A), where parameter values are set as *x*_*c*_ = 20, *b* = 2, *k*_off_ = 1 and *μ* = 10. (E) The global scenario for dependence of the TV on both time scale factor *α* and ratio*k*_on_/*k*_off_, where parameter values are set as in (A).

Compare Fig. 3(A) and Fig. S3(A), in Fig. 3(A), we observe that if ratio *k*_on_/*k*_off_ is fixed, there is an optimal timescale factor such that the MFPT is minimal, but in Fig. S3(A), the MFPT is a monotonically decreasing function of timescale factor if ratio *k*_on_/*k*_off_ is fixed.

